# Functions of TIAM1 at the interface of centriole assembly and autolysosome cycling

**DOI:** 10.64898/2026.07.02.735969

**Authors:** Paula A. Coelho, Chenrong Yu, David M. Glover

## Abstract

Centrosome amplification is frequently associated with chromosomal instability and tumor progression, but how cells coordinate centriole assembly with the control of centrosome numbers and quality remains poorly understood. TIAM1 is a RAC1 guanine nucleotide exchange factor previously implicated in centrosome-associated signaling and βTrCP-dependent control of PLK4 abundance. Here, we examined how Tiam1 regulates autophagy-lysosome homeostasis in mouse embryonic fibroblasts induced to overexpress PLK4. In contrast to a previous model in which Tiam1 loss promotes productive centriole overduplication, we found, by super-resolution imaging and expansion microscopy, an abnormal distribution of PLK4 on the centrioles centriole-associated structures following TIAM1 depletion, suggesting that TIAM1 may support the organization or maturation of centrioles. TIAM1 depletion also resulted in increased LC3B-positive puncta and enlarged LAMP1-positive compartments, but this was not accompanied by increased LC3B-II accumulation after bafilomycin A1 treatment.

These findings suggest that TIAM1 may act at the interface between centriole assembly and endolysosomal/autolysosomal organization, linking TIAM1 to lysosome-associated centrosome quality-control pathways.

## Introduction

Centrosome amplification is frequently observed in cancer cells and has long been associated with chromosomal instability, altered mitotic fidelity, and tumor progression^1–4^. Although extra centrosomes can promote multipolar spindle formation and chromosome segregation errors, cancer cells can also activate adaptive mechanisms that allow them to tolerate or cluster supernumerary centrosomes^2,3^. How cells sense centrosome amplification and restore centrosome homeostasis remains incompletely understood.

We previously identified a RAC1-ATG16L1-dependent autophagy pathway that is activated in response to centrosome amplification and limits the persistence of supernumerary centrosomes^5^. This pathway is regulated by two opposing RAC1 modulators. ARHGEF2 functions as the guanine nucleotide exchange factor that activates RAC1 and promotes autophagy-mediated centrosome clearance, whereas ARHGAP15 acts as a counteracting RAC1 GAP that restrains RAC1-ATG16L1 signaling^5^. Together, ARHGEF2 and ARHGAP15 define a reciprocal signaling module linking cytoskeletal regulation, RAC1 activity, autophagy, and centrosome-number control. These findings raised the broader question of whether additional guanine nucleotide exchange factors contribute to centrosome homeostasis by coordinating mitotic signaling with autophagy-lysosome function.

Several GEFs have previously been implicated in mitotic or centrosome-associated processes. ECT2, a RhoA GEF, is regulated by centrosomal Aurora A and contributes to mitotic signaling^6^, whereas Intersectin-2, Cdc42 GEF, controls spindle orientation during epithelial morphogenesis^7^. Among RAC1 GEFs, TIAM1 provides one of the clearest links between RAC1 signaling and centrosome-associated regulation. TIAM1, or T-lymphoma invasion and metastasis 1, was originally identified as an invasion-inducing gene for T-lymphoma cells^8^. It activates RAC1 and connects upstream signaling pathways to actin cytoskeletal remodeling, migration, invasion, survival, and metastasis^8–10^. In line with these functions, high TIAM1 expression has been associated with lymphatic metastasis and poor prognosis in multiple solid tumours^11^. In mouse cancer models, TIAM1 deficiency delayed Neu/HER2-induced mammary tumor initiation and reduced metastasis, although this effect was not observed in Myc-driven mammary tumors ^12^. TIAM1 has also been linked to chemoresistance and invasive behavior in colorectal cancer, where TIAM1 inhibition increased sensitivity to chemotherapy and reduced invasion^13^.

Beyond its canonical role in RAC1-dependent cytoskeletal regulation, TIAM1 has been implicated in centrosome-associated mitotic signalling. TIAM1 and RAC1 localize to centrosomes during prophase and prometaphase, and loss of TIAM1/RAC activity increases inter-centrosomal distance, suggesting that TIAM1-RAC1 signaling restrains centrosome separation and counterbalances Eg5-driven forces before mitotic entry^14^. TIAM1 also functions upstream of centrosomal PAK activation in prophase^15^. More recently, TIAM1 has been reported to localize to centrosomes during S phase and to regulate centriole duplication by controlling centrosomal PLK4 abundance through a βTrCP-dependent mechanism^16^. In this model, *Tiam1* depletion increases PLK4 levels at the centrosome and elevates CENTRIN3 puncta, interpreted as centrosome overduplication. This phenotype can be rescued by wild-type TIAM1 but not by a TIAM1 mutant unable to bind βTrCP, suggesting that TIAM1 promotes βTrCP-dependent PLK4 degradation. This mechanism is in accord with earlier work showing that SCF/Slimb-βTrCP-mediated turnover limits PLK4/Sak abundance and centriole overduplication^17–19^.

These observations position TIAM1 as a cell-cycle-regulated signaling factor that contributes to centrosome function and mitotic fidelity, rather than as a constitutive structural component of the centrosome. Our identification of an ARHGEF2-RAC1-ATG16L1 pathway controlling autophagy-dependent centrosome clearance^5^ led us to discover that TIAM1 also contributes to centrosome homeostasis through both RAC1-dependent centrosome signaling and a distinct function linked to autophagy and lysosome regulation.

## Results

### Tiam1 depletion does not induce centriole amplification in TetON-Plk4 MEFs

To determine whether TIAM1 contributes to centriole-number control, we depleted TIAM1 in mouse embryonic fibroblasts established from a mouse line in which expression of PLK4 can be driven from a doxycycline-inducible promoter (Tet^ON^-Plk4 MEFs)^5^ and quantified centriole number under basal conditions and following doxycycline-induced Plk4 overexpression (Figure 1A, B). In contrast to the published model in which *Tiam1* depletion promotes PLK4 accumulation and centriole overduplication^16^, we found that reduced TIAM1 levels did not induce centriole amplification under basal conditions. In control MEFs, only 2.42% ± 0.66% of cells exhibited centriole amplification, a frequency not significantly different from *Tiam1*-depleted cells, in which 1.97% ± 0.90% of cells showed amplification (Figure 1B).

**Figure 1.**
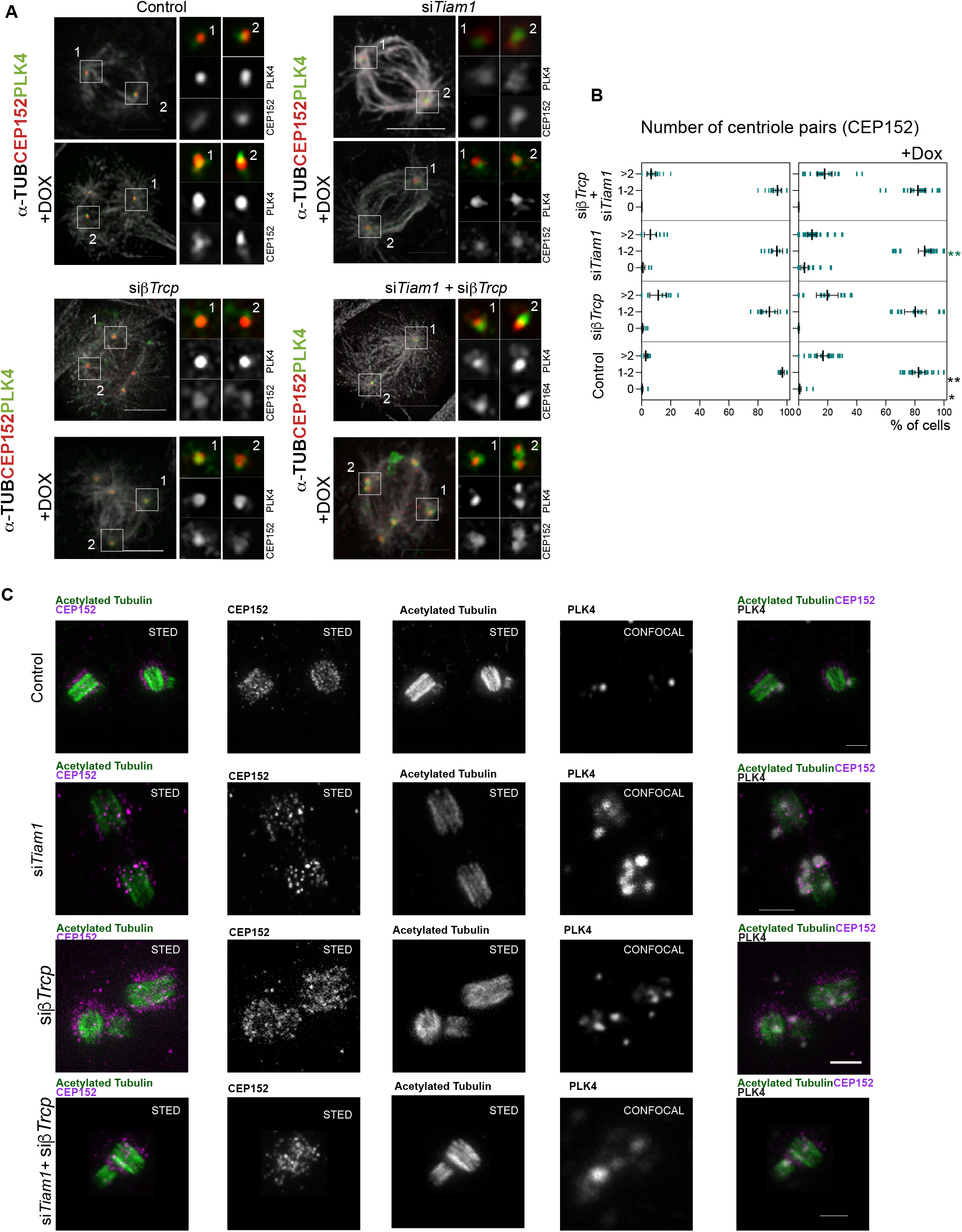
TIAM1 depletion does not induce centriole amplification and does not enhance βTrCP-dependent centriole overduplication. (A) Representative immunofluorescence images showing centrosome phenotypes in TetON-Plk4 mouse embryonic fibroblasts (MEFs) following siRNA-mediated knockdown of *Tiam1*, β*TrCP*, or combined *Tiam1* and β*TrCP*. Cells were treated with the indicated siRNAs in the absence or presence of doxycycline (−/+Dox) to induce Plk4 overexpression and stained for PLK4 (red), CEP152 (green), and alpha-tubulin (white). Scale bar, 10 μm. (B) Quantification of the percentage of cells in each CEP152-positive centriole-pair category: 0, 1-2, or >2 centriole pairs. Data are from 11 independent biological replicates per condition: control, control + Dox, si*Tiam1*, si*Tiam1* + Dox, siβ*TrCP*, siβ*TrCP* + Dox, si*Tiam1* + siβ*TrCP*, and si*Tiam1* + siβ*TrCP* + Dox. Error bars indicate the mean ± 95% confidence interval (CI). Statistical significance was assessed using the Kruskal-Wallis test followed by Dunn’s multiple-comparisons test. Significant p values are indicated in black for the comparison between control −Dox and control +Dox, and in green for comparisons between control and the corresponding knockdown condition under equivalent PLK4-overexpression conditions. ***p < 0.001, **p < 0.005, *p < 0.05. Source data and full statistical analyses are provided in Supplementary File 1. (C) Representative STED and confocal images of control and si*Tiam1*-treated cells immunostained to reveal acetylated tubulin, CEP152, and PLK4. Acetylated tubulin and CEP152 were imaged by STED microscopy to resolve centriolar architecture, whereas the corresponding PLK4 signal was acquired by confocal microscopy in the same cells. Control cells displayed compact acetylated tubulin-positive centrioles with organized CEP152 signal and discrete PLK4 foci. Representative images of cells depleted of *Tiam1*, β*TrCP*, or co-depleted of *Tiam1* and β*TrCP*, stained and imaged as in control.

Following Plk4 overexpression, *Tiam1* depletion also failed to increase centriole amplification. On the contrary, *Tiam1* knockdown reduced the frequency of centriole amplification from 20.77% ± 3.43% in control cells to 13.15% ± 3.68% in *Tiam1*-depleted cells (Figure 1B). These results indicate that, in TetON-Plk4 MEFs, TIAM1 is not required to restrain centriole overduplication. Rather, TIAM1 appears to support the productive formation or maintenance of centriole amplification induced by elevated PLK4.

We then depleted both *Tiam1* and β*TrCP* and found that the combined depletion did not produce the strong increase in productive centriole amplification that would be expected if loss of TIAM1 simply stabilized PLK4 through impaired βTrCP-dependent turnover. Instead, *Tiam1/*β*TrCP* co-depleted cells exhibited predominantly one-to-two centriole-pairs, and the frequency of cells with more than two centriole pairs largely paralleled that observed after β*TrCP* depletion alone (Figure 1B).

Together, these data indicate that TIAM1 is not required to restrain centriole amplification in TetON-Plk4 MEFs but raise the question whether TIAM1 might support the organization or maturation of PLK4-positive centriole assembly intermediates, particularly when PLK4 homeostasis is challenged by β*TrCP* depletion or Plk4 overexpression.

### Tiam1 depletion enhances PLK4 puncta at the centriole

To address the above questions we next combined super-resolution STED microscopy on tissue-expanded specimens, to determine whether *Tiam1* depletion affects centriole architecture or the organization of PLK4 at centrioles (Figure 1C). In control cells, procentrioles were positioned close to the proximal region of the mother centriole, adjacent to regions occupied by CEP152 and associated with discrete PLK4 foci. This organization accords with normal centriole assembly, in which PLK4 is spatially restricted to promote procentriole formation at defined sites. Typically, a single PLK4 *focus* is found at the most proximal region of the parent where the procentriole is observed. By contrast, TIAM1-depleted cells displayed centriole-related structures of abnormal morphology including smaller procentriole-like structures that strikingly, were associated with multiple large foci staining for both PLK4 and its CEP152 partner protein (Figure 1C). High levels of PLk4 were also seen in cells undergoing low levels of centrosome amplification following depletion of βTrCP alone that showed low levels of centriole amplification reflecting the expected increase in PLK4 levels in the absence of SCF function. The double depletion of βTrCP and TIAM1 resulted in centriole of similar organization. Although high levels of PLK4 were observed at the centriole after TIAM1 depletion, this did not lead to centriole amplification suggesting that it represents other functions of TIAM1 in maintaining centriole organization. We conclude that TIAM1 does not directly affect the centriole amplification even though it leads to raised levels of PLK4.

### Tiam1 depletion increases LC3B-positive structures without increasing LC3B-II accumulation after lysosomal inhibition

Because our previous work identified an autophagy-dependent pathway controlling centrosome homeostasis ^5^, we next asked whether TIAM1 depletion affects autophagy or autophagosome-lysosome maturation. We used cells expressing the tandem mCherry-EGFP-LC3B reporter to assess autophagic structures as green fluorescence is lost in the final lysosomal environment. context of Plk4 overexpression (Figure 2A-C). We found that TIAM1-depleted cells displayed increased numbers of LC3B-positive puncta compared with control cells and these were further increased following induction of PLK4. These puncta included LC3B-positive structures located near PLK4/CEP164-positive centrosomal material. The ratio of mcherry to GFP fluorescence indicated the highest levels of autophagy in TIAM1-depleted cells generating amplified centrioles following PLK4 induction. Because an increase in LC3B-positive puncta can reflect either increased autophagosome formation or impaired progression through the autophagy pathway ^20^, we compared LC3B-II levels before and after treatment with bafilomycin A1, which blocks lysosomal acidification to block autophagic flux (Figure 2D,E). TIAM1 depletion did not cause a further increase in LC3B-II accumulation after bafilomycin A1 treatment compared with control cells, suggesting that the increase in LC3B-positive structures observed after TIAM1 depletion is not accompanied by increased autophagic flux. Thus, TIAM1 is likely to be required for autophagosome maturation, trafficking, or autophagosome-lysosome progression.

**Figure 2.**
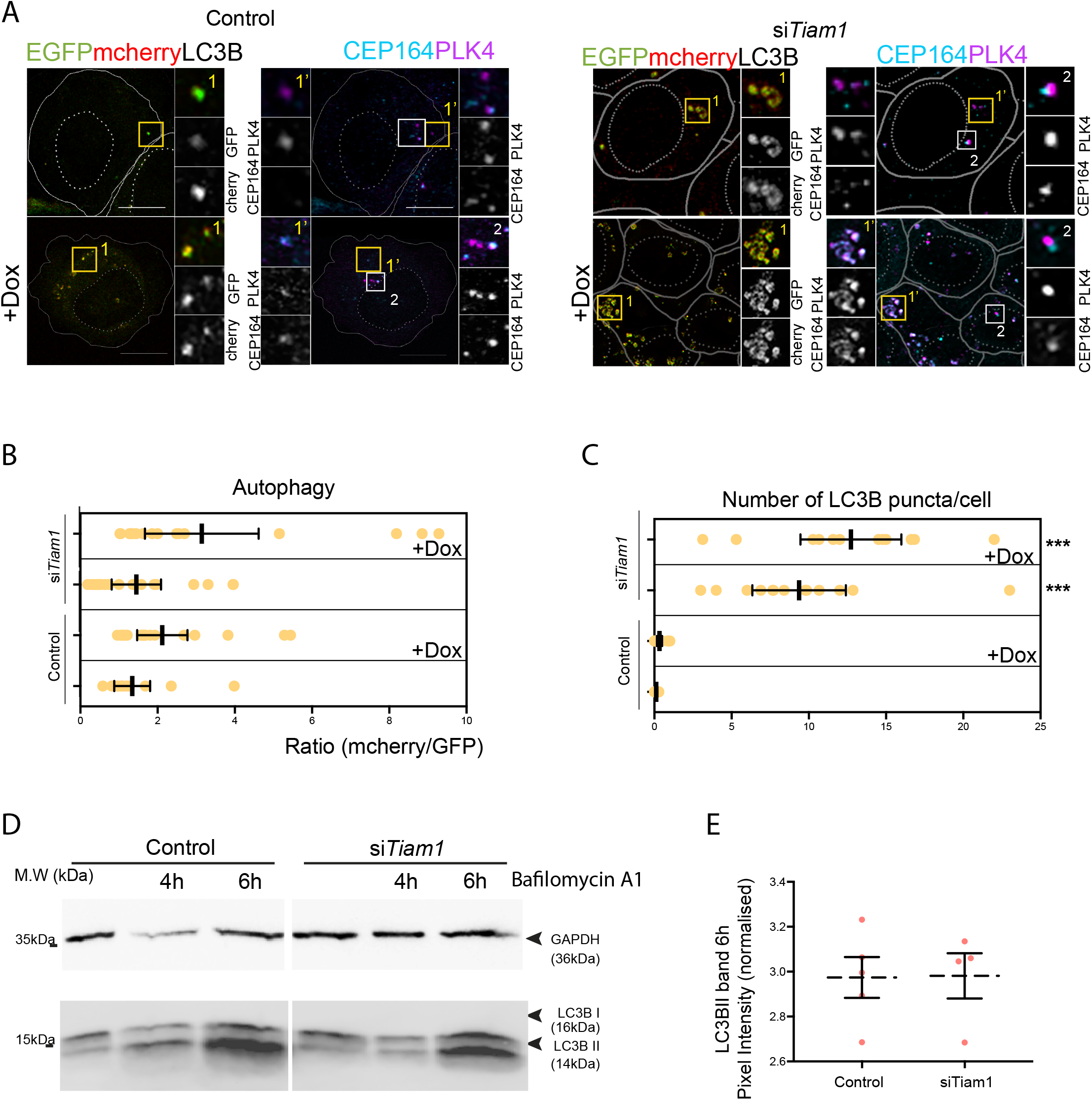
TIAM1 depletion increases LC3B-positive autophagic structures without increasing LC3B-II accumulation after lysosomal inhibition. (A) Representative images of control and si*Tiam1*-treated Tet^ON^-Plk4 MEFs expressing the tandem mCherry-EGFP-LC3B autophagy reporter following doxycycline-induced Plk4 overexpression (+Dox). Cells were immunostained for CEP164 and PLK4. The left panels show mCherry::EGFP::LC3B reporter signal, and the right panels show CEP164-positive centrosomes and PLK4. Insets show merged and single channel mCherry and EGFP reporter signal, together with PLK4 and CEP164 staining. *Tiam1*-depleted cells displayed increased LC3B-positive puncta in proximity to PLK4/CEP164-positive structures compared with control cells. Scale bars, 10 μm. (B) Quantification of autophagic reporter signal, expressed as the mCherry/EGFP LC3B fluorescence ratio, in control and si*Tiam1*-treated cells following Plk4 overexpression. (C) Quantification of the number of LC3B puncta per cell in control and si*Tiam1*-treated MEFs following Plk4 overexpression. Statistical analysis was performed using the Kruskal-Wallis test followed by Dunn’s multiple comparisons test; p < 0.0001. (D) Representative immunoblot analysis of LC3B-I and LC3B-II levels in control and si*Tiam1*-treated cells incubated with bafilomycin A1 for 4 or 6 h. GAPDH was used as a loading control. (E) Quantification of LC3B-II band intensity after 6 h of bafilomycin A1 treatment, normalized to GAPDH. Each point represents one independent biological replicate; horizontal bars indicate mean ± SEM. No significant difference was detected between control and si*Tiam1*-treated samples. n = 3 independent biological replicates. Statistical analysis is shown in Supplementary file1.

### Tiam1 depletion perturbs LAMP1-positive endolysosomal and autolysosomal compartments

We next examined LAMP1-positive compartments to determine whether *Tiam1* depletion affects lysosomal or autolysosomal organization (Figure 3A). LAMP1 labels late endosomes, lysosomes, and autolysosomes; therefore, enlarged LAMP1-positive structures do not necessarily indicate increased autophagy or enhanced lysosomal degradation^21^. Nevertheless, LAMP1 staining provides a useful readout of endolysosomal and autolysosomal compartment organization.

**Figure 3.**
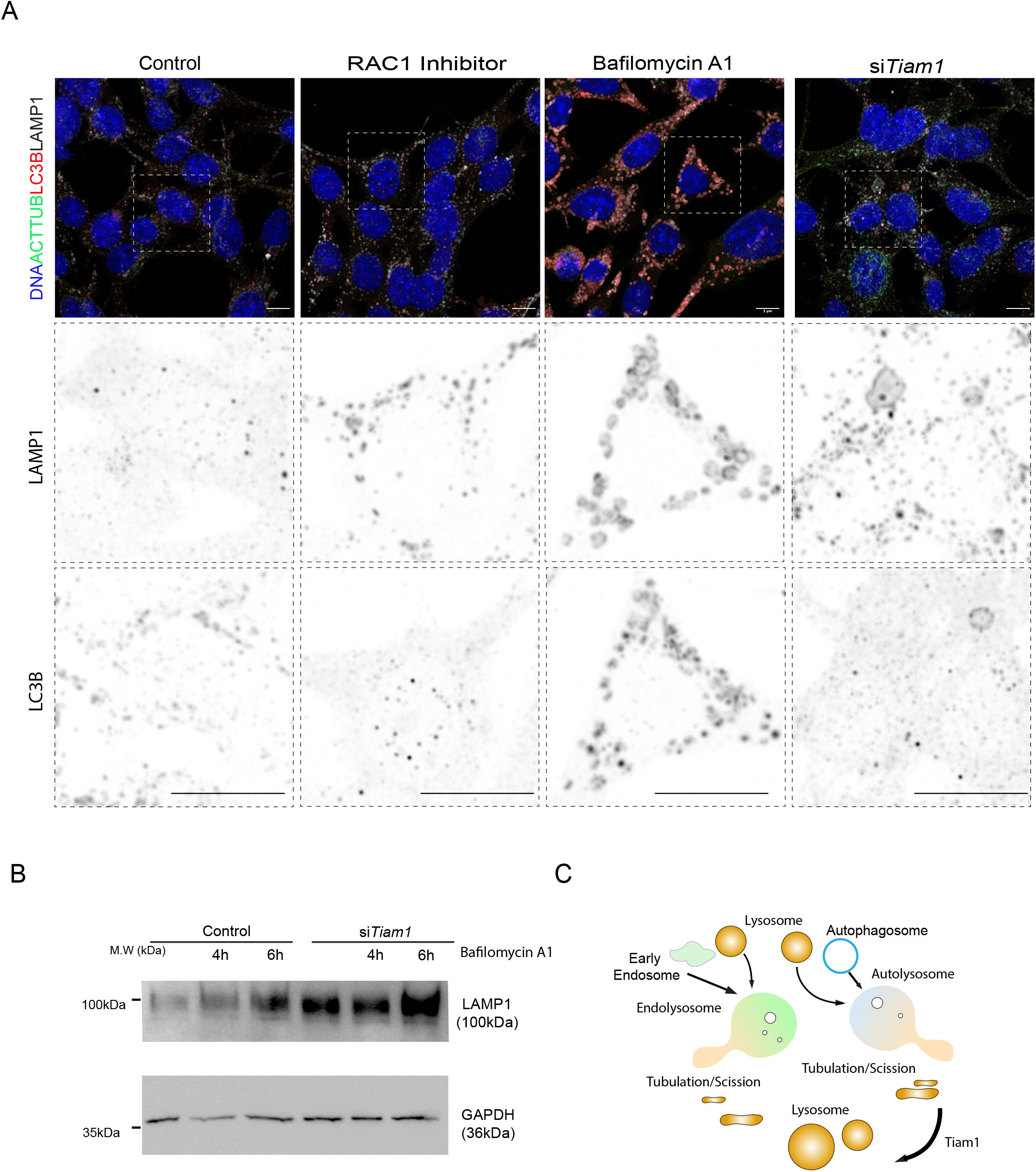
TIAM1 depletion alters LAMP1-positive lysosomal compartments. (A) Representative immunofluorescence images of control cells and cells treated with the RAC1 inhibitor EHT1864, bafilomycin A1, or si*Tiam1*. Cells were stained for DNA (blue), acetylated tubulin (green), LC3B (red), and LAMP1 (white). Split channels for LAMP1 and LC3B are shown below each image (shown as inverted grayscale images). Compared with control cells, si*Tiam1*-treated cells displayed enlarged and/or more prominent LAMP1-positive compartments, consistent with altered lysosomal organization. Scale bars, 10 μm. (B) Representative immunoblot analysis of LAMP1 levels in control and si*Tiam1*-treated cells incubated with bafilomycin A1 for 4 or 6 h. GAPDH was used as a loading control. LAMP1 levels were increased in si*Tiam1*-treated samples but did not further increase with prolonged bafilomycin A1 incubation, suggesting that *Tiam1* depletion alters the abundance or organization of LAMP1-positive compartments without inducing additional bafilomycin-dependent LAMP1 accumulation. (C) Working model showing the proposed role of TIAM1 in lysosome and autolysosome maturation. TIAM1 is proposed to contribute to endolysosomal and/or autolysosomal tubulation or scission, thereby supporting lysosome recycling from endolysosomal and autolysosomal compartments.

*Tiam1*-depleted cells displayed enlarged and more prominent LAMP1-positive compartments compared with control cells (Figure 3A). These structures were also observed in cells stained for LC3B, supporting the idea that *Tiam1* depletion alters the morphology of autophagy-lysosome compartments. Importantly, treatment with the RAC1 inhibitor EHT1864 ^22^ did not reproduce the increase in size and number of LAMP1-positive or LC3B-positive compartments observed after *Tiam1* depletion. This suggests that TIAM1 regulates lysosomal organization through a mechanism that is not simply explained by loss of catalytic activity of RAC1.

Consistent with the imaging data, LAMP1 protein levels were increased in *Tiam1*-depleted cells compared with control extracts (Figure 3B). However, LAMP1 levels did not further increase with prolonged bafilomycin A1 treatment. Together with the LC3B-II data, this indicates that *Tiam1* depletion expands or alters LAMP1-positive endolysosomal/autolysosomal compartments without producing a corresponding increase in bafilomycin-dependent autophagic flux. The enlarged LAMP1-positive structures are therefore more consistent with altered lysosome or autolysosome maturation, trafficking, degradation, or lysosome reformation, rather than enhanced autophagosome production. These data support a model in which Tiam1 affects the organization or maturation of endolysosomal/autolysosomal compartments.

TIAM1 depletion from MEFs does not increase centriole numbers. Instead, it produces abnormal PLK4/CEP152-positive centriole-related structures and enlarged LAMP1-positive compartments. This suggests that TIAM1 contributes to coordinating centriole assembly, centrosome quality control, and endolysosomal/autolysosomal maturation. The precise role of TIAM1 in organizing centriole assembly requires further study but our observations strongly suggest that following deplation of TIAM1, the final assembly of the centriole is disorganized such that aggregates of CEP152 and PLK4 can appear at the centrosome but these do not drive centriole duplication.

TIAM1 appears also to contribute to lysosome and autolysosome maturation. The expansion of LAMP1-positive compartments, together with the lack of any increased bafilomycin-dependent LC3B-II accumulation, suggests a defect downstream of autophagosome formation, potentially at the level of autophagosome-lysosome maturation, lysosomal trafficking, degradation, or lysosome reformation. These observations lead us to propose that TIAM1 may support endolysosomal or autolysosomal tubulation and scission, allowing lysosome recycling from hybrid degradative compartments, a process related to autophagic lysosome reformation^23^. Loss of TIAM1 would therefore lead to enlarged LAMP1-positive structures without necessarily increasing autophagic flux (Figure 3C).

This interpretation is consistent with a broader view of TIAM1 as a cell-cycle-regulated signaling and scaffolding factor that coordinates cytoskeletal dynamics, centrosome function, and membrane trafficking. Although TIAM1 is best known as a RAC1 GEF, its effects on PLK4 organization and lysosomal compartment morphology may involve RAC1-independent scaffolding functions, consistent with previous reports that TIAM1 can regulate centrosomal PLK4 abundance through βTrCP-dependent mechanisms independently of its catalytic RAC1-GEF activity^16^.

We therefore propose that TIAM1 contributes to centrosome homeostasis by coordinating centriole homeostasis with autophagosome-lysosome maturation. This mechanism links mitotic centrosome regulation to membrane trafficking and lysosomal adaptation. We speculate that this could provide a potential route by which cancer cells manage aberrant centrosome content while maintaining their proliferative capacity.

## Methods

### siRNA-mediated gene depletion in MEFs

For siRNA-mediated gene knockdown, MEFs were transfected in a 24-well format using Lipofectamine RNAiMAX. For each well, 6 pmol siRNA duplex were diluted in 100 μL Opti-MEM (Gibco, #31985062) and mixed with 1 μL Lipofectamine RNAiMAX (Invitrogen, #13778075). siRNAs are listed in Supplementary file2. Transfection mixtures were gently mixed and incubated for 20 min at room temperature. MEFs were then diluted to 4 × 10□ cells/mL in complete DMEM growth medium, and 500 μL of the cell suspension was added to each well and mixed with the siRNA transfection mixture. Doxycycline was added 24 h after transfection to a final concentration of 4 μg/mL and refreshed 48 h later. Cells were counted and fixed 96 h after siRNA transfection.

### Immunoblotting

Cells were scraped from dishes in 4× Laemmli sample buffer (Bio-Rad, #1610747). After total protein quantification, equal amounts of protein were subjected to electrophoresis on 7% or 15% Tris-Glycine acrylamide gels (Bio-Rad, 30% acrylamide/bis 29:1, #1610156), transferred to nitrocellulose membranes (Bio-Rad, #1704150), and immunoblotted with the indicated primary and secondary antibodies (Supplementary File 1).

### Immunofluorescence

Cells were grown on coverslips, washed with 1x PBS after removal of the medium, and fixed in cold methanol at -20 °C for at least 12 min. Cells were rehydrated in 1x PBS, followed by permeabilization in PBS containing 0.5% Triton X-100 for 15 min and three washes for 10 min each in PBS containing 0.1% Triton X-100. Blocking was performed in PBS containing 0.1% Tween-20 and 10% FBS for 1 h, followed by incubation with primary antibodies overnight at 4 °C and secondary antibodies for 1 h at room temperature in PBS containing 0.1% Tween-20 and 10% FBS. Washes were performed using PBS containing 0.1% Tween-20. Coverslips were mounted in Vectashield Mounting Medium with DAPI (Vector Laboratories, H1200-10). Images were collected on a Leica SP8 Stellaris with 63×/1.4 oil objectives using Leica Application Suite X software (LAS-X). Images were deconvolved with Huygens Professional version 19.04 (Scientific Volume Imaging, The Netherlands); processing and analysis were performed with ImageJ version 1.53a and Adobe Illustrator 2024. All images shown are projections of all z optical sections, acquired with a z-step of 0.5 μm.

### STED microscopy

Two-dimensional STED images were acquired using an Abberior STEDYCON microscope (Abberior Instruments GmbH, Göttingen, Germany) equipped with a high-numerical-aperture oil-immersion objective (100×, NA 1.45). Images were acquired using STEDYCON software in sequential acquisition mode, with standard confocal and STED settings appropriate for the fluorophores used. For centriole imaging, acetylated tubulin and CEP152 were detected using STED-compatible fluorophores, STAR ORANGE and STAR RED, respectively, and acquired in 2D STED mode. The PLK4 signal was acquired in confocal mode in the same cells using the 488-nm channel. Imaging settings, including laser power, detector gain, pixel size, scan speed, and averaging, were kept constant between experimental conditions within each experiment.

Images were processed in Fiji/ImageJ using identical linear brightness and contrast adjustments for all images within each comparison. No nonlinear image processing was applied. Centriole length and width were measured from 2D STED images in Fiji/ImageJ using acetylated tubulin-positive centriolar structures. STED microscopy was combined with expansion microscopy adapted from the protocol provided by Dr. Meng-Fu Bryan Tsou and Dr. Kanako Ozaki at MSK.

### Statistics and reproducibility

Experiments were performed with at least three independent biological replicates, unless otherwise stated in the figure legends. For microscopy-based quantifications, at least 50 cells were quantified per biological replicate from multiple randomly selected microscopy fields. The total number of cells quantified for each condition is indicated in the corresponding source data. To quantify centriole number revealed by CEP152 staining, cells were scored from randomly selected microscopy fields and assigned to the indicated categories: 0, 1-2, or >2 centriole pairs. The percentage of cells in each category was calculated for each replicate and is shown in the source data. Where indicated, individual microscopy fields are shown in the graphical representation of the data; statistical analyses were performed on biological replicate-level values unless otherwise stated. No data were excluded from the analyses. Statistical analyses were performed using the Kruskal-Wallis H test to assess differences among groups. When significant differences were detected, post-hoc pairwise comparisons were performed using Dunn’s multiple comparisons test with Bonferroni correction. All statistical tests were two-sided, unless indicated otherwise in figure legends. Exact n values, statistical tests and P values are provided in the corresponding source data.

### Graphical representation of centriole number

Individual microscopy samples are represented as bars, with each bar showing the percentage of cells within one of three categories: 0 centrioles, 1–2 centrioles, or >2 centrioles per cell. Data were plotted using GraphPad Prism version 10.6.0. Data represent at least three independent biological replicates, unless otherwise stated in the figure legends. The mean is shown with a black bar, and error bars represent the mean ± 95% confidence interval (CI).

### LC3B and LAMP1 western blotting for quantifying autophagic flux

Cells were treated with bafilomycin A1 (Baf A1; Cell Signaling Technology, #50-205-0494) at 1 μM for 4 and 6 h prior to sample collection. Cells were washed with ice-cold PBS and lysed in RIPA buffer containing protease and phosphatase inhibitors. Western blotting was performed using membranes sequentially incubated with antibodies to detect LC3B, LAMP1 and GAPDH.

Band intensities were quantified using Image Lab v6.1.0 (Bio-Rad) from ChemiDoc MP images and normalized to GAPDH from the same membranes. Individual data points are provided in the Supplementary File 1 indicated. Quantifications were performed from three independent biological replicates. Two-group comparisons were analyzed using the Mann-Whitney U test.

